# Mechanosensitive junction remodelling promotes robust epithelial morphogenesis

**DOI:** 10.1101/648980

**Authors:** Michael F. Staddon, Kate E. Cavanaugh, Edwin M. Munro, Margaret L. Gardel, Shiladitya Banerjee

## Abstract

Morphogenesis of epithelial tissues requires tight spatiotemporal coordination of cell shape changes. In vivo, many tissue-scale shape changes are driven by pulsatile contractions of intercellular junctions, which are rectified to produce irreversible deformations. The functional role of this pulsatory ratchet and its mechanistic basis remain unknown. Here we combine theory and biophysical experiments to show that mechanosensitive tension remodelling of epithelial cell junctions promotes robust epithelial shape changes via ratcheting. Using optogenetic control of actomyosin contractility, we find that epithelial junctions show elastic behaviour under low contractile stress, returning to their original lengths after contraction, but undergo irreversible deformation under higher magnitudes of contractile stress. Existing vertex-based models for the epithelium are unable to capture these results, with cell junctions displaying purely elastic or fluid-like behaviours, depending on the choice of model parameters. To describe the experimental results, we propose a modified vertex model with two essential ingredients for junction mechanics: thresholded tension remodelling and continuous strain relaxation. First, a critical strain threshold for tension remodelling triggers irreversible junction length changes for sufficiently strong contractions, making the system robust to small fluctuations in contractile activity. Second, continuous strain relaxation allows for mechanical memory removal, enabling frequency-dependent modulation of cell shape changes via mechanical ratcheting. Taken together, the combination of mechanosensitive tension remodelling and junctional strain relaxation provides a robust mechanism for large-scale morphogenesis.

## I. INTRODUCTION

During tissue morphogenesis and repair, individual cells dynamically alter their size and shapes in a highly coordinated fashion, resulting in tissue-scale deformations [1–3]. At the single-cell level, morphogenetic forces are actively generated by dynamic actin filaments in association with myosin-II motors [4]. To effect cell shape changes, active contractions generated by myosin motors must overcome both adhesion forces at cell-cell interfaces, and viscous drag from the environment [5, 6]. *In vivo*, pulses of myosin have been reported to drive cell shape changes in a cyclic fashion as a mechanical ratchet, with cell-cell junctions undergoing a series of contraction, stabilisation, and relaxation [7]. These pulses coordinate robust, tissue-scale deformations during apical constriction [1], where cell apical areas successively shrink to drive bending of the epithelial tissue, or during *Drosophila* germband extension [2], where cell-cell junctions contract and intercalate to elongate the tissue. While myosin-driven contraction pulses are ubiquitous in morphogenetic events, the functional roles of the amplitude or the frequency of contractions remain unknown.

From a mechanistic perspective, tissue cells must be capable of resisting large deformations to maintain their mechanical integrity, while being able to dissipate stresses to undergo remodelling. It remains poorly understood how cells modulate their elastic and viscous properties to respond and adapt to mechanical stresses. A number of different experimental techniques and model systems have been used to probe the viscoelastic properties of epithelial cells. Rheological studies on suspended epithelial monolayers [8] have shown that stress dissipation in strained tissues is controlled by cell divisions, or actomyosin turnover [9, 10]. Optical tweezers have been used to probe the mechanical response of cell-cell junctions under short time scale forces, where they respond elastically [6]. At longer time scales, junctions undergo permanent length changes in response to myosin generated forces [11]. Despite the growing evidence for active remodelling of junction mechanical properties [12, 13], most theoretical models for epithelial mechanics, such as the vertex model [14–16], cellular Potts model [17, 18], or continuum models [19–21], assume constant interfacial tensions and cell contractility. While some recent models attempt to model junctional mechanics [9, 22– 25], these have not been directly tested in experiments at the scale of individual cell junctions.

In this study, we combine theory and biophysical experiments to propose a new model for epithelial cell shape control via mechanosensitive remodeling of junctional tension and strain. Using optogenetic control of RhoA [26], the upstream regulator of actomyosin contractility, we study how adherens junction length responds to acute tension changes of varied amplitude and duration. We find that epithelial cell junctions behave elastically in response to short timescale activations of RhoA. Under longer timescale activation of RhoA, junction contraction eventually stalls and does not recover to its initial length upon RhoA removal. The existing vertex-based models for epithelial mechanics are unable to capture these results, with junctions displaying either purely viscous or elastic responses, regardless of the magnitude of applied stress. We thus propose a new model, in which junctional tension remodels irreversibly only when deformed above a threshold value. In addition, junctions undergo continuous strain relaxation to allow for the removal of mechanical memory. Taken together, thresholded tension remodelling and continuous strain relaxation captures the mechanical behaviour of intercellular junctions under stress, and predicts mechanical ratcheting under episodic activations of contractility, in quantitative agreement with experimental data. Our model provides a potential new understanding of pulsatile contractions that have been widely observed during morphogenesis in vivo [1, 7]. In particular, pulsatile contraction enables epithelial junction shortening further beyond the limit of a single prolonged contraction pulse.

## II. MECHANICAL BEHAVIOR OF EPITHELIAL CELL JUNCTIONS

### A. Vertex model predictions

To develop a mechanistic understanding of junctional length regulation in response to applied mechanical stresses, we first implemented a vertex-based model [27], which has been widely used to model epithelial tissue remodeling [24, 28], wound healing [15, 29, 30], collective motility [31], tissue growth [32, 33], and cell-matrix adhesions [34]. In the commonly used vertex models for two-dimensional tissues [14, 27, 33], the geometry of each cell is defined by a polygon, with cell-cell junctions represented by linear edges and three way junctions by vertices (Fig. 1a). The tissue mechanical energy is given by:

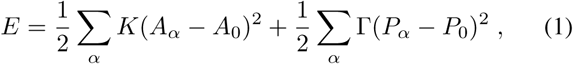

where the first term represents 3D incompressibility of cells with *K* as the height elastic constant, *A*_*α*_ is the planar area of cell *α*, and *A*_0_ is the preferred area. The second term in Eq. (1) results from a combination of actomyosin contractility in the cell cortex and intercellular adhesions, where Γ is the elastic constant for contractility, *P*_*α*_ is the perimeter of cell *α*, and *P*_0_ is the preferred perimeter. The geometric parameters of the vertex model define a target shape index 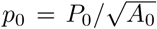, which controls the mechanical properties of the tissue [14, 28]. In particular, the tissue behaves like an elastic solid for *p*_0_ < 3.81, while it flows like a viscous liquid for *p*_0_ > 3.81 [14].

**FIG. 1.**
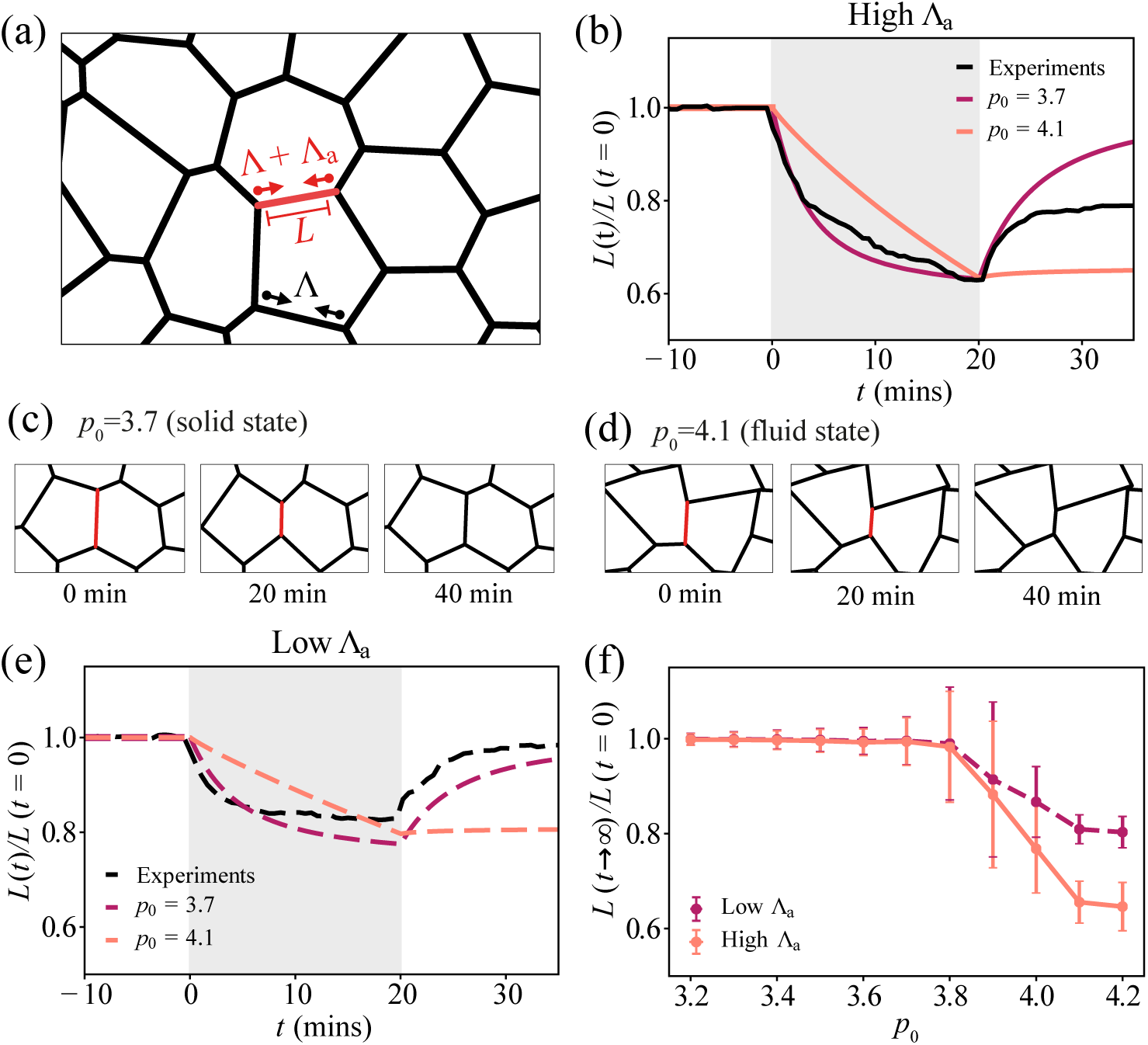
Elastic and viscous response of intercellular junctions in the vertex model. (a) Schematic image of the vertex model. The red edge represents the junction under applied tension, Λ_*a*_. (b) Normalised junction length over time, during and after a 20-minute activation for high applied tension. Grey shaded region indicates the activation period. Red curve: *p*_0_ = 3.7, orange curve: *p*_0_ = 4.1, black curve: experimental data for 20-min activation of RhoA using optogenetics, averaged over 6 edges. (c-d) Simulation images of a junction before, during, and after a 20-minute activation, for (c) a solid tissue, *p*_0_ = 3.7, and (d) a fluid tissue, *p*_0_ = 4.1. (e) Normalised junction length over time, during and after a 20-minute activation for high applied tension. (f) Normalised final junction length against preferred shape index *p*_0_, for high and low values of applied tension, Λ_*a*_. For each value of *p*_0_, 15 simulations were run. Error bars indicate ± 1 standard deviation. For parameters see Tables I and II.

In the over-damped limit, motion for each vertex *i* is governed by:

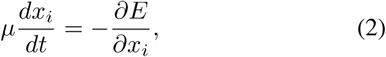

where *µ* is the drag coefficient, and *x*_*i*_ is the position of vertex *i*. In the absence of active or external forces, the tension Λ_*ij*_ on edge *ij*, connecting vertices *i* and *j*, is given by Λ_*ij*_ = *∂E/∂L*_*ij*_, where *L*_*ij*_ is the length of edge *ij*. For the hamiltonian in Eq. (1), Λ_*ij*_ = ∑*_α_* Γ(*P*_*α*_− *P*_0_), where the sum is over all cells containing the edge *ij*. To simulate the effect of time-dependent myosin induced contractions on edge *ij*, we apply an additional tension Λ_*a*_ = Γ_*a*_*L*_*ij*_*/*2, where Γ_*a*_ represents the product of myosin density and the force exerted by each myosin unit [35].

Upon application of myosin contractility over a finite duration (Fig. 1b), the vertex model predicts that cell junctions immediately contract while eventually slowing down for a tissue in the solid state (Fig. 1b-c). After the activation period, the elastic tissue recoils back to its original length, while a fluid tissue exhibits no recoil and remains permanently deformed. In the fluid state (Fig. 1b,d), cells can freely adjust their area and perimeter in response to applied stress, resulting in edges under no tension. A fluid tissue can undergo small junctional deformations with no energy cost, resulting in permanent changes in junction length after applied contraction (Fig. 1d). By contrast, cell junctions are under tension for a solid tissue, stable to small mechanical perturbations. Consequently, contracted edges return to their original lengths after the activation period (Fig. 1b,c).

### B. Comparisons with experimental data

To test the predictions of the vertex model, we use optogenetic control of RhoA in Caco-2 cells [26, 36–38]. By targeting light at chosen cell-cell junctions we are able to increase actomyosin contraction in a highly localised region (Appendix B). Applying a 20-minute activation, we observe a rapid contraction of the junction to ~60% of its initial length (Fig. 1b). After RhoA activation, the junction recoils and recovers to ~80% of its initial length. This indicates a viscoelastic response of the junction, with elastic response on short time scales and viscous behaviour, by permanent length changes, on longer time scales. These data stand in contrast to the predictions of the vertex model. Furthermore, by tuning the light intensity, we are able to control the amount of contractile stress on the junction. When applying 50% of the light intensity, the initial junction contraction rate drops by half. This is followed by a slow phase of contraction that eventually stalls (Fig. 1e). After the activation period, the junction recoils back to its original length, akin to an elastic material.

To simulate low light activation of RhoA, we applied half the active tension Λ_*a*_ in simulations. As before, we find an elastic response of the junction for *p*_0_ < 3.81, and no recoil in the fluid state (Fig. 1e). Testing several other values of the preferred shape index *p*_0_, we observe no deformation for the solid state and permanent tissue deformation in the fluid state, at both high and low values of Λ_*a*_ (Fig. 1f). Thus the vertex model is unable to capture the viscoelastic response of cell junctions under high contractile stress. Our data also stand in contrast to Ref. [11], where cell-cell junctions are modelled as Maxwell viscoelastic elements. Under this model, even low contractile stresses should continuously shorten the junction, in contrast to our experimental data.

## III. JUNCTION LENGTH REMODELLING

Our experimental data show that irreversible junction deformations occur only for sufficiently high amplitudes of contraction, suggesting a thresholded viscoelastic response of intercellular junctions. To this end, we modify the existing vertex model to incorporate irreversible junctional length remodelling above a threshold strain. We adapt the vertex model to treat each cell edge as an elastic spring with spring constant *Y*, and a dynamically changing rest length, 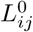 (Fig. 2a). The rest length remodels according to:

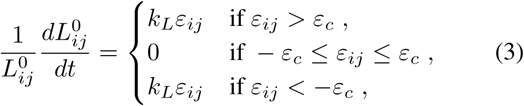

where 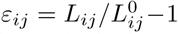 is the strain on edge *ij, ε*_*c*_ is a critical strain beyond which rest length remodelling is triggered, and *k*_*L*_ is the rate of remodelling. The tension on each edge is given by:

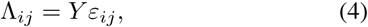

such that the mechanical energy of the tissue is:

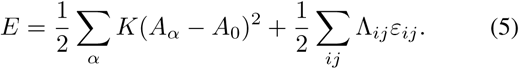

**FIG. 2.**
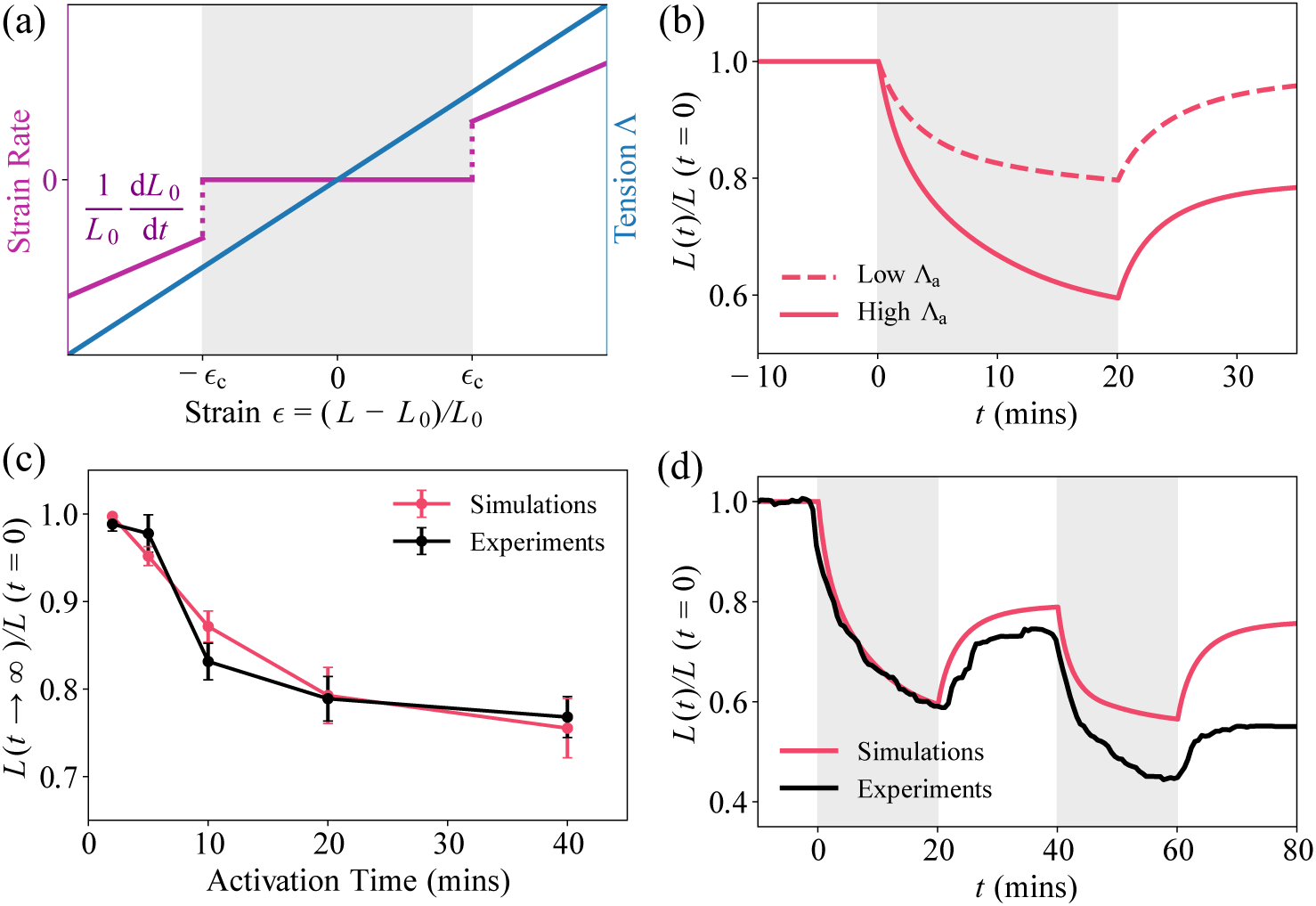
Threshold strain is required to initiate junction remodelling. (a) Rate of change in rest length, 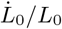, and edge tension Λ, as a function of junction strain *ε*. (b) Model predictions for normalised junction length over time for a 20-minute step pulse of contraction, for high (Λ_*a*_) and low (Λ_*a*_*/*2) values of applied tension. (c) Comparing experimental data and model predictions for (normalised) final junction length vs contraction time. Error bars indicate ±1 standard deviation (Experiments: 2 mins, n=5; 5 mins, n=9; 10 mins, n=3; 20 mins, n=6; 40 mins, n=3. Simulations: all times, n=15). (d) Normalised junction length vs time for two consecutive 20-minute contraction pulses, for experiments (black, n=3) and simulations (red, n=15).

This model predicts that junction contraction is biphasic upon activation of a single contraction pulse of magnitude Λ_*a*_. For high Λ_*a*_, junction length shortens quickly, and then begins to slow down and eventually stall (Fig. 2b). Once the strain in the junction exceeds *ε*_*c*_, the junction rest length begins to remodel and becomes shorter. After the activation period, the junction recoils, but a shorter rest length leads to am increased tension in the edge, resulting in a shorter final length of the junction (Fig. 2b). When applying half the tension Λ_*a*_, the critical strain *ε*_*c*_ is never reached. As a result the rest length remains unchanged and the junction recoils back to its initial length after activation (Fig. 2b). Thus, very short timescale activations are unable to contract the junction beyond the critical strain, leading to elastic response and perfect length recovery (Fig. 2c). Longer timescale activations are required to trigger junctional remodelling. However, remodelling slows down with increasing activation period, thereby limiting the amount of length shortening as observed in experiments (Fig. 2c).

As the final length saturates to 80% of its initial value, irrespective of the magnitude of Λ_*a*_ or the activation period, we investigated if a second contraction pulse could overcome this length saturation. In optogenetic experiments, a second 20-minute activation pulse of RhoA (with a 20-minute rest in between) leads to ratcheted contraction (Fig. 2d). Junction length after the second contraction drops to 80% of its value after the first contraction. However, this behaviour is not reproduced by our model for thresholded junctional length remodelling. After the second activation, the junction length recoils back to its value after the first activation (Fig. 2d).

To understand the mechanistic origin of these results, we compute the edge tension and strain during the pulsatile stress protocol (Fig 3a). During the first activation period, the edge is compressed, resulting in a negative tension acting to expand the junction length. As the strain drops below the critical value, *-ε*_*c*_, the rest length begins to remodel at a rate proportional to the strain. This results in rest length shortening, which brings back the strain to its critical value, *-ε*_*c*_, at which tension and strain remodelling stops. After the removal of exogenous tension, the junction length recoils beyond the rest length, resulting in a positive tension and a permanently strained (shortened) junction (Fig 3a). During the second activation, the junction contraction stalls at the fixed point *ε* = *-ε*_*c*_, and the rest length is unable to remodel further. Since tension is proportional to strain, the critical strain determines the maximum steady state tension possible, Λ_max_ = *Y ε*_*c*_. Above this tension magnitude, the rest length will remodel until strain reaches the critical value. Due to the invariance in Λ_max_, we obtain the same final edge length in the model after any number of activations.

**FIG. 3.**
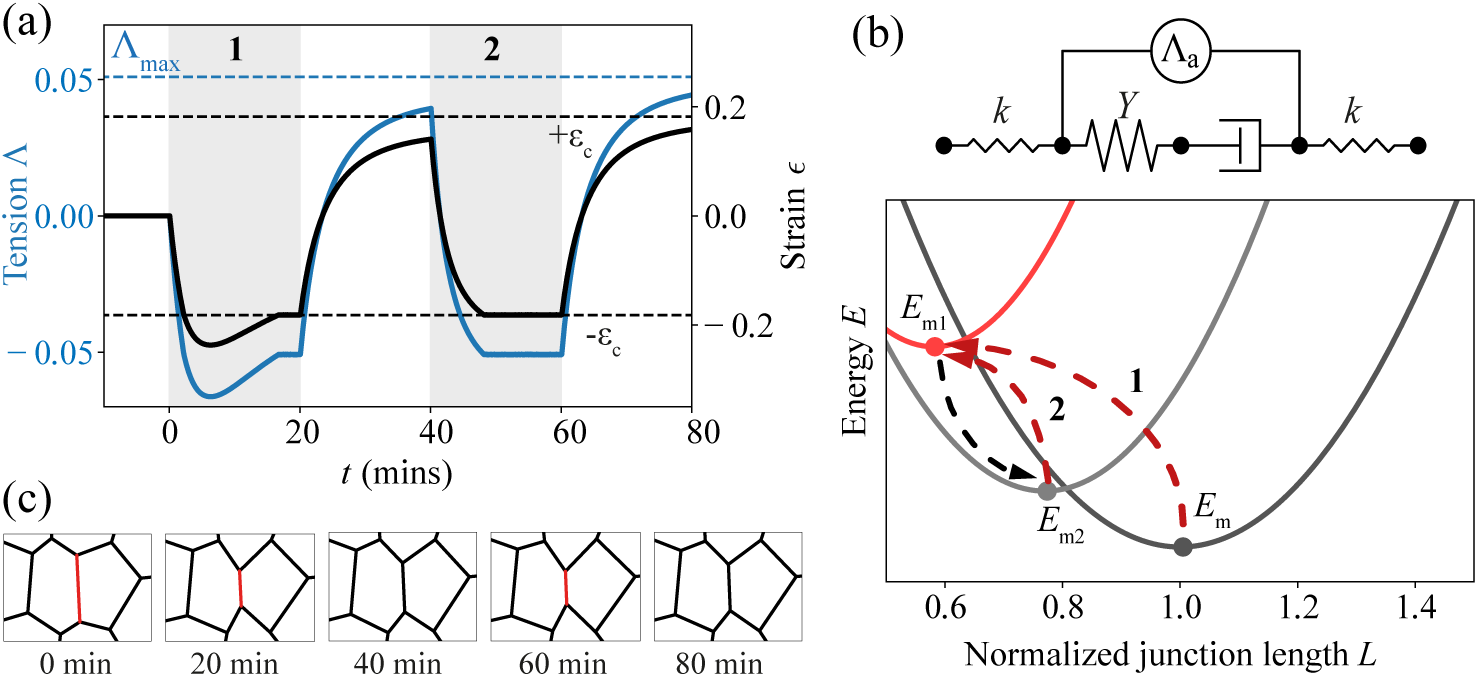
Tension remodelling is essential for ratcheted contraction. (a) Junction tension and strain vs time for two 20-minute activations. The dashed blue line indicates the maximum steady-state tension, Λ_max_ that can be achieved in this model. Dashed black lines indicate the strain threshold. (b) Top: Schematic of the effective medium model. Bottom: Dynamics of energy landscape vs junction length (normalised by its initial length). The arrows indicate the directions of energy shifts during contraction (red) and after contraction pulse removal (gray). The numbers correspond to the pulse number. (c) Simulation images of a junction over time during two consecutive contractions, showing no ratcheting.

### A. Effective medium model

For ratcheted contractions to occur, cell junctions must relax strain to remove mechanical memory, i.e. junctions must overcome their mechanical energy barriers and not get stuck in a local energy minimum. To understand the underlying physics, we constructed an effective medium model for the cell junctions (Fig 3b-Top). Here we treat forces from the neighbouring cells as a spring-like restoring force, while the junction acts like a Maxwell viscoelastic material that requires a threshold strain for the viscous element to remodel. The energy of this effective system is given by:

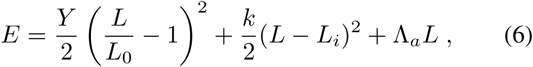

where *L* is the current junction length, *L*_*i*_ is the initial length, and *L*_0_ is the junction rest length. The first term represents the harmonic restoring force from the junction, the second term describes the spring-like restoring force from neighbouring junctions with a spring constant *k*, and the final term represents the applied tension Λ_*a*_ = Γ_*a*_*L/*2. Junction rest length remodels according to Eq. (3). Before the activation of contractility, *L* = *L*_*i*_ = 1, as defined by the minimum of the energy functional, *E*_m_ (Fig 3b).

During the first activation, Λ_*a*_ ≠ 0 and the rest length *L*_0_ shortens. This shifts the energy landscape, which now has a new minimum, *E*_m1_, defining the contracted junction length (Fig 3b)

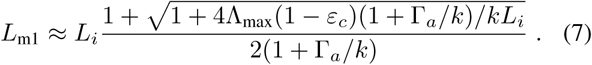

After the first activation, the energy minimum shifts to a higher value, *E*_m2_, due to junction recoil. The new steady state junction length is given by

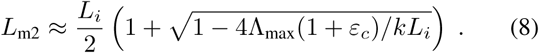

As subsequent contraction pulses are unable to change the rest length, the steady-state length of the junction between the minima (*L*_m1_, *E*_m1_) and (*L*_m2_, *E*_m2_) (Fig 3b). Therefore, ratcheting behaviour is not programmed in this model. For the junction to contract further, an increase in edge tension is required above its maximum possible value, Λ_max_ = *Y ε*_*c*_ (Fig 3a).

## IV. MECHANOSENSITIVE TENSION REMODELLING

In order to capture ratcheted contractions, we developed a model where the edge tension is dynamically remodelled depending on the junctional strain. In this model, the rest length of each edge, 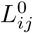, remodels at a rate *k*_*L*_ to match the current junction length (Fig 4a):

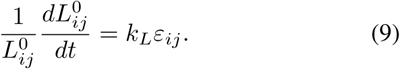

**FIG. 4.**
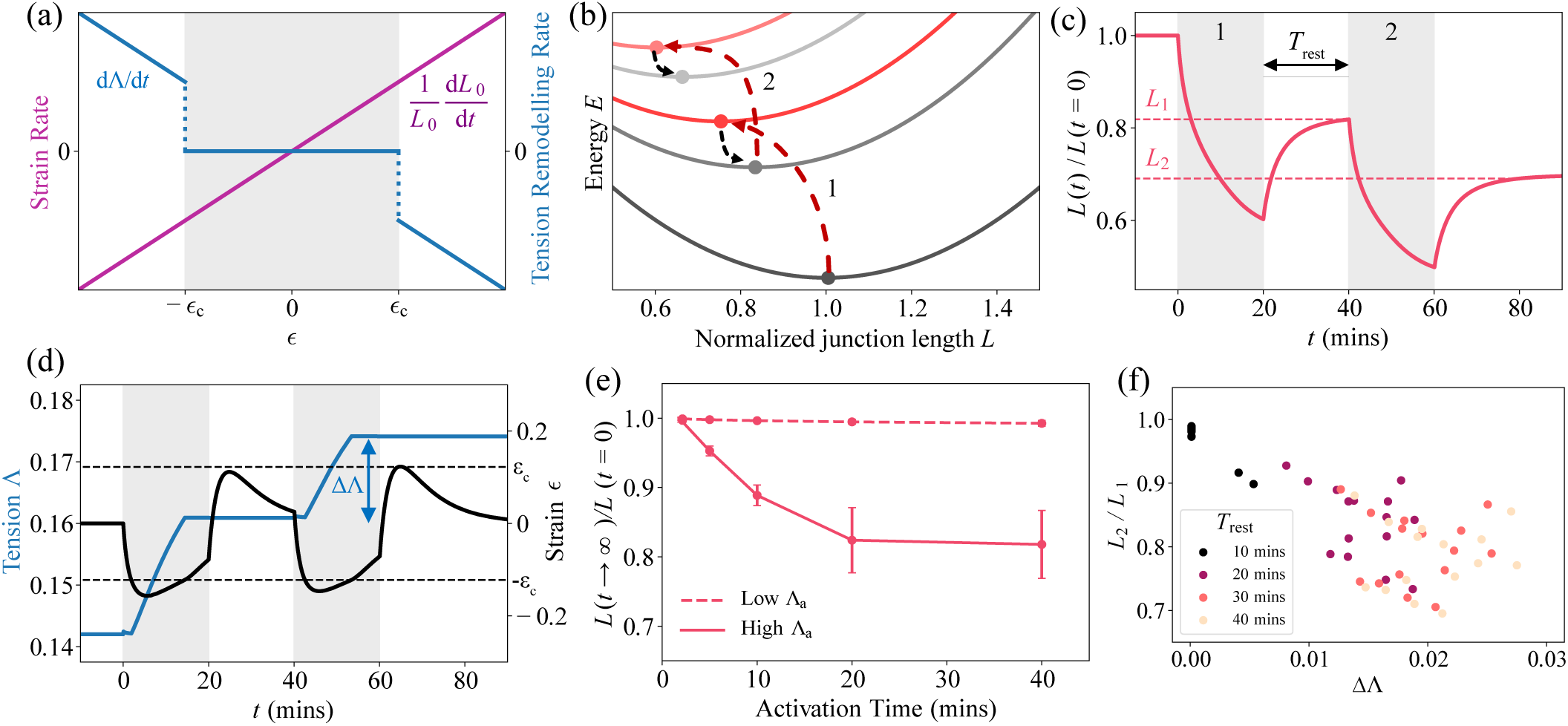
Tension remodelling promotes ratcheted contractions. (a) Model for tension remodelling: rate of change in rest length, 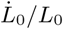, and rate of change in tension, 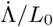, as a function of the junctional strain *ε*. (b) Junction energy landscape in the tension remodelling model, as a function of normalised junction length. (c-d) Dynamics of (c) Normalised junction length, and (d) junction tension and strain, for two successive 20-minute contraction pulses. (e) Normalised junction length vs contraction time, for high and low values of Λ_*a*_. Error bars represent ± 1 standard deviation (n=15). (f) Length ratio *L*_2_*/L*_1_ vs the change in tension ΔΛ between two consecutive contraction pulses, for different values of *T*_rest_. Each point represents a different simulation.

If the strain in the junction exceeds a threshold magnitude, *ε*_*c*_, then the tension begins to remodel:

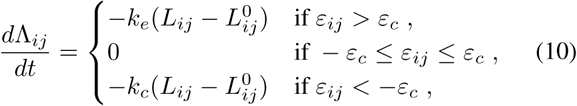

where *k*_*e*_ and *k*_*c*_ are the rates of tension remodelling under junction extension or contraction, respectively (Fig 4a). Thus, if an edge undergoes a large and rapid contraction, the tension will begin to increase and the junction will remain irreversibly shortened.

An effective medium model for this system can be described by the energy function:

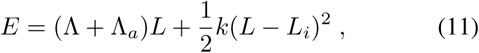

where the first term represents the total tension in the junction, whose dynamics are described by Eq. (10). Applying consecutive pulses of contractions would repeatedly increase the edge tension, and shift the steady-state junction length *L* = (*L*_*i*_ *−* Λ*/k*)*/*(1 + Γ_*a*_*/k*) (given by the minimum of *E*) to successive lower values (Fig 4b). This would result in successive reductions in the final junction length. Note that tension remodelling alone is not adequate to capture the experimental data. Without rest length remodelling, an increased tension would rapidly contract cell edges without limit, leading to their eventual collapse.

### A. Ratchet-like contractions

We find that the combination of continuous strain relaxation [Eq. (9)] and thresholded tension remodelling [Eq. (10)] is sufficient to capture the experimentally observed mechanical behaviour of epithelial junctions, as well as ratcheted contractions upon episodic activation of contractility. In agreement with experimental data (Fig 2d), the junction shrinks to ~ 80% of its initial length after the first contraction, while after the second contraction pulse, the normalised junction length is 70%, roughly 85% of its length after the first pulse (Fig 4c). During each activation, the strain drops below the critical strain and tension is irreversibly increased, allowing the junction length to shrink further than after the first activation (Fig. 4d). Unlike the model with purely rest length remodelling (Fig. 3), combination of tension and rest length remodelling allows the junctional strain to relax back to zero after the removal of exogenous tension (Fig. 4d). Continuous strain relaxation allows for the removal of mechanical memory, which is crucial for promoting contraction below the critical strain during each successive pulse.

As very short timescale activations are unable to trigger tension remodelling, the junctions recoil back to their original length, in line with experimental data (Fig 4e). By contrast, longer activation periods increase the amount of tension remodelling and junction length shortening. For single activations, junction shortening stalls for very long activation periods (>20 minutes), in agreement with experimental data (Fig 2c). Thus ratcheting provides a mechanism to further shorten junctions past the single contraction limit. Applying a lower tension during activation is unable to sufficiently strain the junction, and therefore a combination of high Λ_*a*_ and long activation period is required for junctional remodelling (Fig 4e).

We quantified the efficiency of ratcheting by computing the ratio of lengths before and after the second activation period (Fig 4c). For a given rest period *T*_rest_ between the activations, we define *L*_1_ as the length before the second activation, and *L*_2_ as the length *T*_rest_ minutes after the the second activation. For different values of *T*_rest_, we compare the ratio *L*_2_*/L*_1_ to the change in tension ΔΛ after the second activation (Fig 4d). We find a robust negative correlation between *L*_2_*/L*_1_ and ΔΛ (Fig 4f). A longer *T*_rest_ allows the junctional strain to relax to zero when Λ_*a*_ = 0. Consequently, we obtain a more effective ratchet, with a higher ΔΛ and a lower *L*_2_*/L*_1_ (Fig 4f).

### B. Convergence-extension movements

To test the effectiveness of mechanical ratcheting for tissue-scale deformation, we applied our model to simulate convergent-extension movements. Convergent-extension is a widespread mode of tissue morphogenesis, where a tissue segment converges along one axis as an oscillatory ratchet, while extending along the other axis via cell rearrangements [39]. To simulate convergent-extension, we generated a small tissue of hexagonal cells and applied a time-dependent tension, Λ_*a*_, along its horizontal edges, in five 40-minute pulses (Fig 5a,b).

**FIG. 5.**
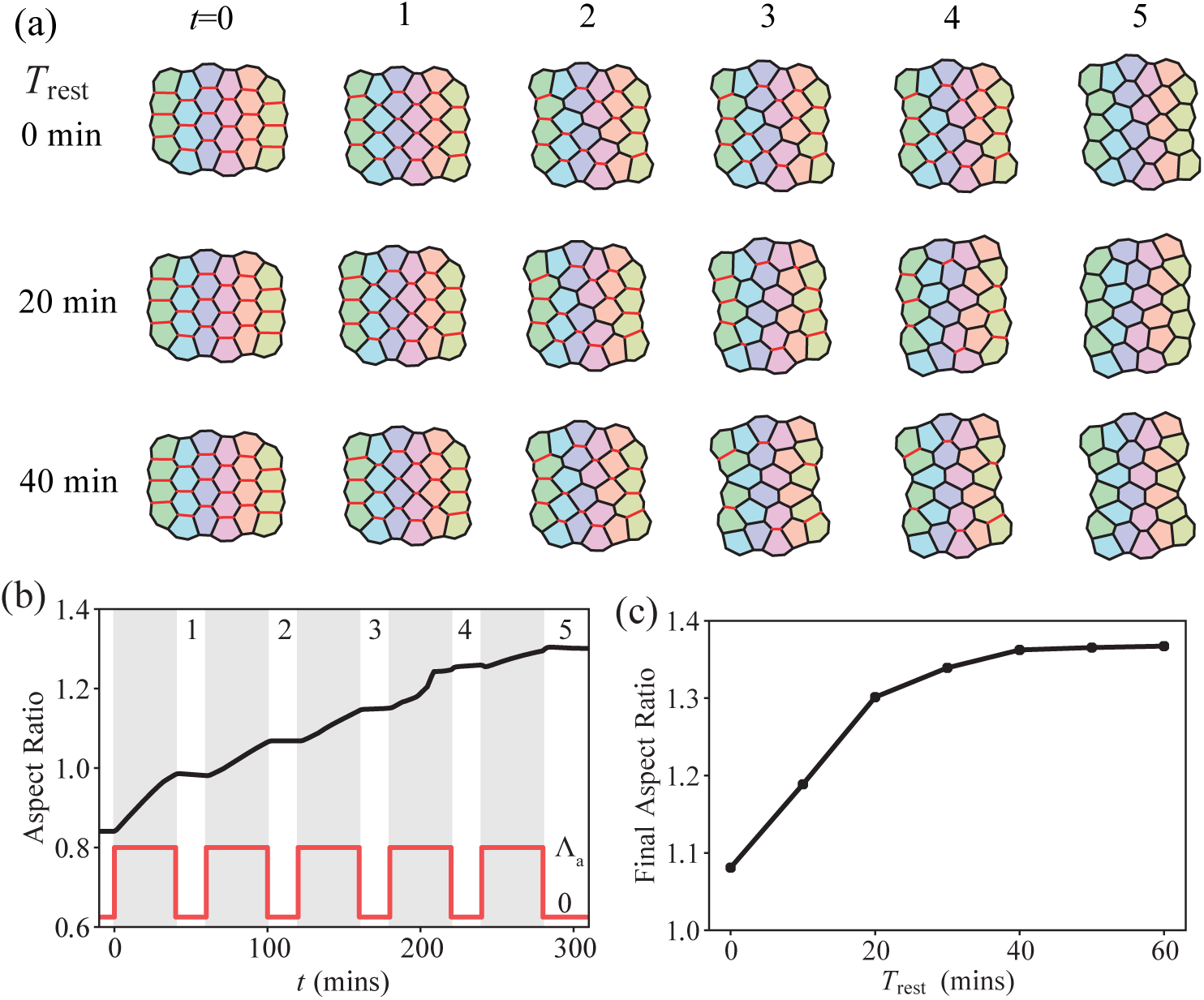
Convergence-extension is more effective for longer rest periods between successive activations. (a) Tissue configurations during five 40-minute contractions at different rest periods (*T*_rest_) between contractions. Numbers 1-5 indicate the number of rest cycles the tissue has undergone, as shown in (b). The cell colours indicate their initial location column-wise, and red edges indicate the initially activated edges. (b) Aspect ratio of the tissue vs time, for five 40-minute activations (bottom inset) with 20 minutes of rest periods in between. The red pulse shows the applied tension vs time during the activation and rest cycles. (c) Final aspect ratio of the tissue vs *T*_rest_, showing that longer rest periods between contractions promote more tissue deformation.

Once a cell edge contracts below a threshold length, *L*_min_, it undergoes an intercalation event resulting in cell neighbour exchanges. During activations of contractility, the tissue converges horizontally, and extends vertically via cell rearrangements, resulting in a higher aspect ratio shape (Fig 5a,b). During the rest period between activations, the tissue remains in a deformed state due to irreversible tension remodelling. By allowing longer rest periods between successive contractions, the amount of tension remodelling increases, resulting in a higher aspect ratio structure (Fig 5c) with more cell rearrangements (Fig 5a). Thus, our model for tension remodelling is sufficient to capture tissue-scale movements via ratcheted contractions, as commonly observed in morphogenesis.

## V. DISCUSSION

Pulsatile regulatory dynamics are recurrent in cells, and enable diverse cellular functions through independent control of pulse frequency, amplitude, and duration [40]. In development, pulses of actomyosin have been shown to coordinate epithelial cell shape changes and mechanical stability [1, 7, 41– 43]. However, the functional roles of actomyosin pulsation and the significance of its temporal structure have remained elusive. The present theoretical study, in combination with biophysical experiments, demonstrates the functional roles of the amplitude and frequency of contraction pulses on epithelial morphogenesis. While high amplitude pulsing triggers irreversible junction deformation via tension remodelling, junction shortening eventually stalls for prolonged activation of actomyosin contractility. However, longer rest periods between successive contraction pulses enable a higher degree of junction shortening via mechanical strain relaxation. We show that frequency-dependent modulation of junction deformation, in combination with cell rearrangements, is sufficient to drive tissue-scale shape changes via convergent-extension movements. These results provide a potential new understanding of the significance of pulsatile contractions, suggesting that high amplitude and low frequency myosin pulsing, separated by longer rest periods, is most effective in driving large-scale epithelial remodelling.

Our proposed theory for mechanosensitive epithelial junction remodelling advances the existing cell-based models of epithelial tissues. The widely used vertex model for epithelial mechanics [27], and its existing variants are unable to capture the experimentally measured length dynamics of epithelial junctions upon time-varying contractions. In the vertex model, junctions either contract like a purely elastic material that is reversible upon stress removal, or junctions continuously shorten their lengths like a fluid even for a low amount of applied contraction. These predictions are inconsistent with experimental data, where junctions display an elastic behaviour only under short, or weak activation of RhoA. Longer periods of RhoA activation lead to permanent junction shortening, which saturates for prolonged pulse duration.

To capture the experimental data, we propose a modified vertex model with two essential features of epithelial junction mechanics: thresholded tension remodelling and continuous strain relaxation. In our model, junctions undergo permanent tension remodelling only above a critical strain threshold. This enables irreversible junction deformation for sufficiently strong and prolonged activation of contractility, in agreement with our experimental data. Furthermore, thresholded tension remodelling acts as a filter for small amplitude and frequency fluctuations in contractile activity, imparting mechanical robustness to the system. However, tension remodelling alone is not sufficient to promote large-scale deformations, as junction shortening saturates for prolonged activation of contractility. To this end, continuous strain relaxation in epithelial junctions allows the system to lose memory of its mechanical deformation. As a result, pulsatile activation of contractility, with sufficiently long periods of relaxation, enables large-scale irreversible deformations in epithelia. Taken together, the combination of mechanosensitive tension remodelling and strain relaxation in epithelial junctions provides a mechanical basis for robust morphogenesis in epithelial tissues.

## APPENDIX A: SIMULATION METHODS

### Model Implementation

We non-dimensionalize force scales by 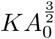 and length scales by 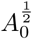, setting *K* = 1 and *A*_0_ = 1. Thus, Eq. 1 becomes:

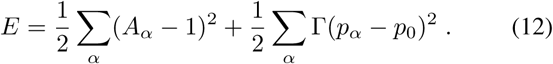

The model is simulated using Surface Evolver [44]. A round tissue of 50 cells is first relaxed without remodelling so that the simulations begin at mechanical equilibrium. 15 different cell edges are randomly selected for activation. After each activation, the tissue is reset to the equilibrium state and then the next cell edge is activated. Any cell edge that contracts below a critical length, 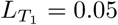, and would decrease in length over the next time step, undergoes a *T*_1_ transition. Here the cells rearrange and the contracted edge is replaced by a perpendicular edge of the same length, following which the edge tension is set to the initial value.

### Model parameters

In the vertex model (Section II), for all values of *p*_0_, the value of contractility, Γ, is taken from Farhadifar et al [33], and we use a constant value for friction coefficient, *µ* (Table I). For each value of *p*_0_, we choose Γ_*a*_(*p*_0_) (Table II) such that the normalized junction length after a 20-minute activation is the same as in the experiments (Fig 1b).

**TABLE I.** Default vertex model parameters

| Parameter | Value |
| --- | --- |
| Contractility, $\Gamma$ | 0.04 |
| Friction coefficient, $\mu$ | 0.25 min |

**TABLE II.** Applied tension for different *p*_0_

| $p_0$ | $\Gamma_a$ |
| --- | --- |
| 3.2 | 0.0708 |
| 3.3 | 0.0603 |
| 3.4 | 0.0497 |
| 3.5 | 0.0392 |
| 3.6 | 0.0288 |
| 3.7 | 0.0187 |
| 3.8 | 0.0108 |
| 3.9 | 0.0069 |
| 4.0 | 0.0055 |
| 4.1 | 0.0053 |
| 4.2 | 0.0052 |

Parameters for the model in Section III (Table III) and Section IV (Table IV) are fit to experiments by minimizing the mean square difference between the normalized junction length in simulations and the mean length in experiments for 2, 5, 10, 20, and 40-minute activations. We compute the fitting error as:

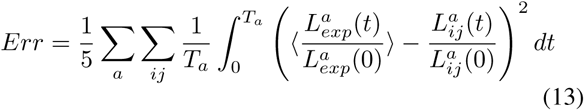

where *a* is the index for activation time (*a* ∈ *{*2, 5, 10, 20, 40*}*), *T*_*a*_ is the total experimental time, *ij* indicates the activated junction in the simulation, and 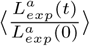 is the normalized junction length averaged over all experiments.

**TABLE III.** Elastic model parameters

| Parameter | Value |
| --- | --- |
| Rest length remodelling rate, $k_L$ | $0.227 \text{ min}^{-1}$ |
| Critical strain, $\epsilon_c$ | 0.091 |
| Spring constant, $Y$ | 0.280 |
| Applied contractility, $\Gamma_a$ | 0.337 |
| Friction, $\mu$ | 5.502 min |

**TABLE IV.** Tension remodelling parameters

| Parameter | Value |
| --- | --- |
| Rest length remodelling rate, $k_L$ | $0.159 \text{ min}^{-1}$ |
| Critical strain, $\epsilon_c$ | 0.122 |
| Initial tension, $Y$ | 0.142 |
| tension remodelling rate, $k_c$ | $0.020 \text{ min}^{-1}$ |
| tension remodelling rate, $k_e$ | $0.000 \text{ min}^{-1}$ |
| Applied contractility, $\Gamma_a$ | 0.085 |
| Friction, $\mu$ | 1.132 min |

## APPENDIX B: OPTOGENETIC EXPERIMENTS

Cells stably expressing the TULIP optogenetic system [38] were plated atop polymerized collagen gels coating a 4-well chamber (Ibidi) at a final concentration of 2mg/ml. Cells were plated at least two days before optogenetic experiments to ensure a polarized and mature epithelial monolayer. Cell-cell junctions were delineated with CellMask Deep Red plasma membrane stain (Molecular Probes, Life Technologies). Junctions were illuminated by a 405nm laser for 1000 ms before each image acquisition every 35 seconds. Regions were drawn in Metamorph (Molecular Devices, Sunnyvale, CA) and manually adjusted in real time to ensure the blue light was restricted to the activated junction. Junction lengths were manually measured in FIJI software using the segmented line tool. A Mosaic digital micromirror device (Andor) was used for optogenetic recruitment using a 405nm laser. A Nikon Ti-E (Nikon, Melville, NY) with a Yokogawa CSU-X confocal scanning head (Yokogawa Electric, Tokyo, Japan) and laser merge model with 491, 561, and 642nm laser lines (Spectral Applied Research, Ontario, Canada) was used to image cells. The objective used was a 60x 1.49 NA ApoTIRF oil immersion objective (Nikon) or a 60x 1.2 Plan Apo water (Nikon) objective.

## ACKNOWLEDGEMENTS

MFS is supported by EPSRC (UK Engineering and Physical Sciences Research Council) funded PhD studentship. SB acknowledges support from the Royal Society (URF/R1/180187). KC is supported by an HHMI Gilliam Fellowship, National Academies of Sciences Ford Foundation Fellowship, and NIH training grant (GM007183). MLG acknowledges funding from NIH RO1 GM104032. EM acknowledges funding from NIH RO1 HD099931.

